# Skeletal Metastasis of Prostate Cancer Is Augmented by Activation of EphA2 Noncanonical Signaling and Ligand-Deficient Bone Microenvironment

**DOI:** 10.1101/2024.09.25.615079

**Authors:** Ryan Lingerak, Aaron Petty, Xiaojun Shi, Soyeon Kim, Hebei Lin, Jer-Tsong Hsieh, Dean Tang, Bangyan Stiles, David Wald, Sooryanarayana Varambally, Arul M. Chinnaiyan, Zhenghong Lee, Colm Morrissey, Bingcheng Wang

## Abstract

EphA2, a member of Eph family receptor tyrosine kinases (RTKs), is overexpressed in multiple types of solid human tumors, particularly at the late stages. However, whether and how it drives specific malignant processes remain elusive. We report that EphA2 is elevated during prostate cancer (PCa) progression in multiple syngeneic murine models. Interestingly, human metastatic PCa specimens from two Rapid Autopsy Programs showed selective overexpression of EphA2 in metastasis to the bone, but not to the lymph nodes or viscera. Serine 897 phosphorylation that mediates the pro-oncogenic, noncanonical signaling by EphA2 was also upregulated in bone metastasis. Analysis of human datasets shows EphA2 overexpression is associated with skeletal but not visceral metastases. Ephrin-A1, a major cognate ligand for EphA2, is lost in PCa bone metastasis, which is correlated with poor prognosis. Further, the bone microenvironment is unique in expressing little of the five *EFNA* genes, providing permissive microenvironment for bone colonization. S897A mutation that ablates EphA2 noncanonical signaling, suppressed PCa development. Restoration of ephrin-A1 expression in PC-3, a model cell line for double negative prostate cancer derived from bone metastasis and devoid of ephrin-As, profoundly changed global tyrosine phosphorylation profiles, inhibited basal ERK and Src activities in vitro, and suppressed tumor development in the bone. Together these results demonstrate EphA2 overexpression and concomitant loss of ligands in PCa lead to activation of noncanonical signaling that is sustained in ephrin-A1-deficient skeletal milieu to promote bone metastasis.

## Introduction

Prostate cancer (PCa) is the most common non-skin cancer in men and is the second leading cause of cancer death in men within the United States.^1^ While usually indolent or benign, a small fraction (5%) rapidly progress to malignant diseases^2,3^. PCa is initially responsive to androgen deprivation therapy (ADT) or castration. However, aggressive forms of the disease inevitably become resistant to this therapy^4,5,6^, leading progressively to metastatic castration resistant PCa (mCRPC), a fraction of which further progress to neuroendocrine prostate cancer (NEPC) and double negative PCa (DNPC) where both AR and NEPC markers are lost. Prostate cancer frequently metastasizes to bone with 70%–90% of patients having radiographically detectable skeletal involvement. Clinically, bone metastases remain an incurable and contribute significantly to patient morbidity and mortality. Discovery of molecular and cellular mechanisms behind PCa progression to skeletal metastasis is imperative to identifying new therapeutic targets and areas of intervention.^7^

EphA2 receptor tyrosine kinase (RTK), a member of the Eph subfamily of RTKs, has been previously implicated in PCa. EphA2 is overexpressed in mCRPC, but not in early localized PCa tumors^8^, which was later confirmed by others^6,9^. However, the molecular mechanisms and pathological significance of EphA2 in mCRPC remain poorly understood. We have reported previously that EphA2 has dual opposing roles as both an oncogene and a tumor suppressor, and can switch between the roles dependent on binding of its ephrin-A ligands.^10^ Ligand stimulation leads to EphA2 canonical signaling characterized by tyrosine kinase catalytic activation and inhibition of Ras/ERK^11^, PI3K/Akt ^10^ and β1 integrin-mediated signaling pathways,^12^ pointing to tumor suppressive role. Genetic studies confirmed this notion, as EphA2 knockout mice displayed dramatically increased susceptibility to carcinogenesis.^13^ In the absence of ligand, EphA2 becomes a target of AGC family serine/threonine kinases including Akt, RSK and PKA that phosphorylate serine 897 (pS897), an event that converts Epha2 from a tumor suppressor into an oncogenic proteins that promotes tumor cell migration, stemness, and development of drug resistance in multiple cancer types.^10,14,15^ Molecularly, recent studies using the time-resolved live cell spectroscopy revealed that the EphA2 noncanonical signaling is enabled by the head-tail interaction between the ligand binding domain and the second fibronectin type II repeat of two adjacent receptors.^16^

Interestingly, Morrissey et al. show that ephrin-A1 was one of the top three genes lost in mCRPC, especially in bone metastases.^17^ The loss of ligand expression and gain of receptor expression tilt the balance EphA2 signaling toward the pro-oncogenic, noncanonical signaling and function of pS897-EphA2. However, whether EphA2 noncanonical signaling through pS897 is activated during PCa progression toward mCRPC in human diseases has yet to be documented, and nor is it known if it contributes to metastatic PCa progression in vivo including its preferential bone colonization.

To fill this major gap, we interrogated the relationship of EphA2 signaling and the malignant progression of PCa. Using a combination of cell and animal models, public datasets, as well as human tissue samples from two Rapid Autopsy Programs, we found that EphA2 is dynamically upregulated during PCa progression to mCRPC concomitant with increased serine 897 phosphorylation, and loss of ephrin-A1 ligand. Importorantly, we report EphA2 overexpression and pS897 signal in human bone metastatic specimens, demonstrating activation of EphA2 ligand-independent pro-oncogenic signaling, which was sustained in the bone milieu largely devoid of any ephrin-A ligand expression. Lastly, we show that PCa tumor growth is diminished by either inhibition of non-canonical signaling with the S897A mutation, and restoration of EphA2 expression in PC-3 cels, a model for DNPC originally derived from bone metastasis and devoid of endogenous ephrin-A expression, blocked xenograft growth in the bone. Together our data establish a role of EphA2 in progression toward mCRPC wherein the increased EphA2 expression and decreased ligand enables abundance of ligand-independent signaling via phosphorylation of EphA2,^9^ which results in proliferation, migration, and metastasis of PCa.

## Materials and Methods

### Gene Expression Analysis from Publicly Available Datasets

Expression of *EPHA* receptors and *EFNA* ligands as well as expression of *AR*, and *KLK3* by human PCa cell lines was presented as log_2_ transformed fragments per kilobase of transcript per million fragments mapped (FPKM). Gene expression information was obtained from a dataset accumulated by Smith R. et al. *Scientific Reports* **2020**. These data utilize human cell lines characterized as being castration resistant, androgen sensitive, or having an intermediate phenotype. Bulk RNA sequencing was performed as described in Smith R. et al. *Scientific Reports* **2020.**

Data showing correlations of *EPHA receptor* expression to *AR* expression were acquired from 208 human metastatic PCa biopsies as part of the SU2C/PCF Dream Team published in *PNAS* **2019**. Data points in the correlation plots represent mRNA expression z-scores relative to all samples. Statistical analyses of Pearson correlations presented were performed with GraphPad Prism 9. The mRNA expression dataset was accessed through cBioportal.^18,19^

Patient survival Kaplan Meier plots were generated using data from both the SU2C/PCF Dream Team (*PNAS*, **2019**.) and MSK (*Cancer Cell*, **2010**). datasets. The SU2C/PCF Dream Team dataset consists of at least 96.8 percent metastatic tissue samples, while the MSK dataset consists of 75.4 percent primary prostate tumor samples. In the SU2C/PCF data set *EPHA2* mRNA expression is reported as z-scores relative to all samples. Our EphA2 “High” expressing group consisted of expression scores 1-2.5 and 0-0.25 and our EphA2 “Low” consisted of -1.75-0. In the MSK data set both *EFNA1* and *EPHA2* were separated into expression quartiles and the top 25 percent expressing samples were considered the “High” group, while all remaining samples were considered as “Low” expression. Logrank tests and plotting were performed with GraphPad Prism 9. The datasets were accessed through cBioportal.^18,19^

EphrinA expression from non-tumor human tissues was exported from the Human Protein Atlas^20^ and analyzed with GraphPad Prism 9 (Human Protein Atlas, https://www.proteinatlas.org/). Data presented consist of data from GTEx represented as normalized transcripts per million log_10_. Data normalization was performed previously by the Human Protein Atlas.

### Tissue Microarray

Human specimens used to confirm a possible link between bone metastasis and EphA2 upregulation, and to investigate phosphorylation of EphA2 at S897 in bone metastases, we collaborated with the Pacific Northwest Prostate Cancer SPORE through Dr. Beatrice Knudsen, and Dr. Colm Morrissey. Of significant interest to us, they have constructed tissue microarray (TMA) from the bone and soft tissue metastases of PCa as part of the SPORE’s Rapid Autopsy Program. Rigorous quality control and patient documentation are in place for the tissue procurement. TMAs have been verified as suitable for IHC detection of multiple phospho-proteins, requisite for the studies proposed in this application.

### Cell Lines and Reagents

PC-3, DU145, LNCap, CWR22R, and C4-2 cell lines were obtained from the American Type Culture Collection (ATCC, Manassas, VA). These cells were cultured in RPMI 1640 media supplemented with 10% fetal bovine serum, 10 mg/ml glutamine, 100 U/ml penicillin, and 0.1 mg/ml streptomycin. LAPC4 and LACP9 cells were obtained from Dean Tang with permission from Robert Reiter (UCLA) and cultured in Iscove’s Modified DMEM supplemented with 10% fetal bovine serum, 10 mg/ml glutamine, 100 U/ml penicillin, and 0.1 mg/ml streptomycin with the addition of 1nM DHT. TRAMPC1 cells were obtained as a gift from Norman Greenberg and were cultured in DMEM supplemented with 5% fetal bovine serum 5% Nu serum, 10 mg/ml glutamine, 100 U/ml penicillin, and 0.1 mg/ml streptomycin with the addition of 10 nM dehydroepiandrosterone (DHEA). MPC3 and MycCaP cells were cultured in DMEM supplemented with 10% fetal bovine serum, 10 mg/ml glutamine, 100 U/ml penicillin, and 0.1 mg/ml streptomycin. 293FT cells used in production of lentiviral constructs were obtained from Life Technologies (Grand Island, NY) and cultured in DMEM media supplemented as above, but with addition of 0.1 mM sodium pyruvate, 1 mM non-essential amino acids (NEAA), and 500 μg/ml G418. Cells were maintained in a humidified incubator at 37° C and 5% CO_2_ . All cell lines were routinely tested for mycoplasma contamination. Additional mycoplasma screens were performed 1-2 days prior to injection into mice.

Ephrin-A1-Fc was produced as described previously.^12^ Rabbit anti-pEphA/B antibody was raised against the conserved phosphopeptides in the juxtamembrane regions (Miao et al. 2009).^10^ Rabbit polyclonal pS897-EphA2 antibody was produced as described previously^10^, as well as purchased (Cell Signaling Technology #6347S). Rabbit polyclonal anti-ephrin-A1 antibody was purchased from Santa Cruz Biotechnology (sc-911). Other antibodies were purchased as follows: Rabbit monoclonal anti-GAPDH (Cell Signaling Technology #2118S), rabbit monoclonal anti-pS473-Akt (Cell Signaling Technology #4060), rabbit polyclonal anti-pErk1/pErk2 (Cell Signaling Technology #9101S), rabbit monoclonal anti-EphA2 (Cell Signaling Technology #6697S), rat monoclonal anti-EphA2-PE (R&D Systems FAB639P), rabbit monoclonal anti-AR (Abcam ab133273), rabbit polyclonal anti-pY772-EphA2 (Cell Signaling Technology #8244S), rabbit monoclonal anti-pY588-EphA2 (Cell Signaling Technology #12677S), mouse monoclonal anti-Tubulin (R&D Systems MAB9344)

Goat anti-rabbit and goat anti-mouse secondary antibodies conjugated to horseradish peroxidase (HRP) were purchased from Santa Cruz Biotechnology (Santa Cruz, CA) and from Biorad (#1706515, and #1706516 respectively). For immunohistochemistry, mouse monoclonal anti-EphA2 was purchased from Millipore (Bedford, MA). Mouse monoclonal anti-tubulin was purchased from Sigma (Saint Louis, MO). Donkey anti-rabbit RedX conjugated secondary antibody was purchased from R&D Systems (Minneapolis, MN).

### Cell Lysate Immunoblotting

Cells were cultured as described and lysate was collected when cells were subconfluent (∼70% confluent). Media and cells were lysed in modified RIPA Buffer as described previously,^12^ and stored in loading buffer containing sodium dodecyl sulfate (SDS) and phosphatase inhibitors. Lysates were separated using the Bolt System (Life Technologies, Grand Island, NY) and 4-12% Bis-Tris Plus gradient gels, followed by transfer to PVDF membranes according to manufacturer’s instructions. Membranes were blocked for 1 hour at room temperature in 3% BSA in 0.05% TBS-T, followed by overnight incubation in appropriate primary antibodies. Cells were then washed in 0.05% TBS-T and incubated 1 hour at room temperature in appropriate secondary antibodies, followed by further washing and development with Luminol Reagent (Santa Cruz Biotechnology, Santa Cruz, CA).

### Cell Stimulations

Stimulations with ephrinA1-Fc ligand (EA1 or EA1-Fc) were done in 6 well corning cell culture plates following cell growth for 2 days, or until plates were approximately 70% confluent. Attached cells were treated with 2 μg/ml EA1-Fc for 0, 15 and 60 minutes without changing cell culture media. For stimulations of PC-3 cells with FBS followed a 24 hour period of serum starvation, cells were exposed to either 10% FBS or 10% FBS plus EA1-Fc for a period of 1 hour prior to lysate collection.

### Immunofluorescence Staining

For immunofluorescence staining, cells were cultured on 10 mm square coverslips for 2 days, then stained exactly as described previously.^12^ Briefly, cells were fixed in 4% paraformaldehyde, followed by permeabilization with 0.5% NP-40 and blocking with Blocking Buffer (3% BSA, 2% goat serum, PBS). Cells were then washed in PBS, then 0.1% BSA/PBS and incubated with rabbit polyclonal anti-EphA2 antibody. Cells were then incubated with RedX-conjugated donkey anti-rabbit secondary antibody, followed by mounting in Mounting Medium with DAPI (Vector Laboratories, Burlingame, CA). Staining was analyzed with a Leica Microsystems microscope and imaged using Metamorph software (Leica Microsystems, Wetzlar, Germany).

### Animal Models and Subcutaneous Injections

All syngeneic animal experiments were completed with C57Bl/6J mice purchased from Jax (Jackson Lab, 000664). All xenograft models were completed with nude mice purchased from Case Western Reserve University. All animal husbandry and procedures were approved by the Institutional Animal Care and Use Committee (IACUC) at Case Western Reserve University. Subcutaneous injections were completed by resuspension of cells in endotoxin-free PBS and injection of both hind flanks of the mouse with 100 μl solution. Cell suspensions were stored on ice during injections for no longer than 1 hour. All animals were monitored 24 hours post-injection as well as twice a week post evidence of tumor formation.

### Serial *In Vivo* Passage of Murine PCa Cells in Syngeneic Host

Parental TRAMPC1 cells were injected subcutaneously into syngeneic C57Bl/6J mice at 5.0 × 10^6^ cells per flank. Post 80 days of tumor formation tumors were excised and dissociated in Collagenase IV containing 10 μM Y27632 dihydrochloride. Collagenase IV was purchased from Sigma-Aldrich (#C4-22-1G), and Y27632 was purchased from AdooQ Bioscience (#A21448). Approximately 0.5 g of tumor tissue was processed with the Milteny OctoDissociater for 41 minutes. Digested samples were immediately passed through a 70 μM cell strainer and washed with 8 ml of DMEM containing 10% FBS. Cells were plated at different concentrations on corning cell culture plates under treatment with fungicide and tetracycline to prevent contamination. Treatment was stopped after 2 days if cells had no bacterial or fungal contamination. After up to 5 passages *ex vivo* and at least 1 freeze thaw cycle, the cells were checked for mycoplasma contamination, and then injected into syngeneic hosts subcutaneously at 5.0 × 10^6^ cells per flank. These injected cells were termed “*in vivo* 1,” (TRAMPC1 IV1) as they had been through a single round of *in vivo* tumor growth.

MycCaP cells were injected subcutaneously into syngeneic FVB mice at 0.7 × 10^6^ cells per flank and monitored for a period of 21 days. Following this, the tumors were harvested and digested for isolation of a cell culture as described above for the TRAMPC1 cells.

MPC3 cells were injected subcutaneously into syngeneic C57Bl/6J mice at 2.0 × 10^6^ cells per flank. Resultant tumors were excised at day 33 post injection. Tumors were digested and cells were isolated as described for TRAMPC1. These resultant MPC3 IV1 cells were checked for mycoplasma and injected again into syngeneic hosts at 2.0 × 10^6^ cells per flank. The resultant tumors were again excised and a new cell line was cultured that had been through 2 rounds of *in vivo* tumor formation called “*in vivo* 2” (MPC3 IV2).

### Live Cell Staining and Cell Sorting

MPC3 cell populations from every time point during the process of *in vivo* selection in syngeneic host (Parent, IV1, IV2), were subjected to live cell staining of EphA2 in order to determine ectopic expression level. Staining was completed with EphA2-PE (anti-mEphA2 PE conjugated R&D Systems #FAB639P). Flow cytometry was completed using the Sony MA900. The mean fluorescence intensity was recorded for each iteration for a minimum of 50,000 cells. This staining took place within the first two weeks of culture post isolation from tumors, excluding the parental MPC3 cells. Each iteration had undergone at least one freeze-thaw cycle.

Parental TRAMPC1 cells were sorted based on level of EphA2 expression also using EphA2-PE antibody. These cells were subjected to fluorescence-activated cell sorting (FACS) using the Sony MA900 sorter which allowed isolation of 30% of the cells with the highest EphA2 expression, and 30% of cells with the lowest EphA2 expression.

### Xenografts Models

PC-3 and C4-2 cells were xenografted subcutaneously onto nude mice in the hind flanks. Cells were prepared in Corning 10 cm tissue culture plates until approximately 70% confluent. Cells were counted, washed with PBS, and mixed with PBS (tested to be endotoxin free). 1.3 × 10^6^ cells were injected into each flank and monitored for a period of 50 days.

### Intratibial Injection

PC-3 cells expressing firefly luciferase and either a vector control, or lenti EphrinA1 expression vector were delivered intratibially using a 28g, ½ inch needle under the patella, through the middle of patellar ligament, and into the anterior intercondylar area in top of the tibia. 2.0 ×10^5^ cells were delivered in 20 μl of PBS at an approximate rate of 5 μl/second. Upon imaging mice were administered 200 μl of 1.25 mg/ml D-Luciferin by intraperitoneal injection (IP) and anesthetized 5 minutes post injection with 1-5 percent isoflurane then imaged using the IVIS Spectrum Bioluminescence Imager. At Week 6, the tibia with PC-3 cell injection were also imaged by planar X-ray to examine bone losses.

## Results

### EphA2 is overexpressed in mCRPC cell lines in vitro and in bone metastasis specimens of human patients in vivo

To explore the pathological roles of EphA2 in the late stage metastatic PCa, we first analyzed gene expression from public bulk RNA sequencing data of human PCa cell lines^22^, and discovered that castration resistant cell lines exhibited overexpression of EphA2 compared to androgen sensitive cell lines, which is consistent with earlier reports of increased EphA2 expression in human metastatic castration resistant specimens (mCRPC)^23^ **(Fig. 1A)**. This is inversely correlated with expression status of AR and KLK3, a downstream target of AR encoding prostate specific antigen (PSA). Immunoblot analysis confirmed the low levels of EphA2 expression in androgen sensitive cell lines, whereas it is upregulated in metastatic castration resistant cells, particularly PC-3 cells derived from bone metastasis that have recently become a model for late stage DNPC **(Fig. 1B)**. This finding was further supported by immunofluorescence staining of EphA2, where it was readily detected in castration resistant DU145 and PC-3 cells, but weakly detected in androgen sensitive LNCaP and CWR22R cells **(Fig. 1C)**. The androgen sensitive LAPC4 are an exception and displays moderate mRNA and protein expression (**Fig. 1A, B)**. Serine 897 phosphorylation (pS897) that demarcates activation of the oncogenic noncanonical signaling is upregulated in metastatic PC3 and DU145 cells, but is absent from androgen sensitive cells, including LAPC4 cells despite moderate EphA2 expression.

**Fig. 1.**
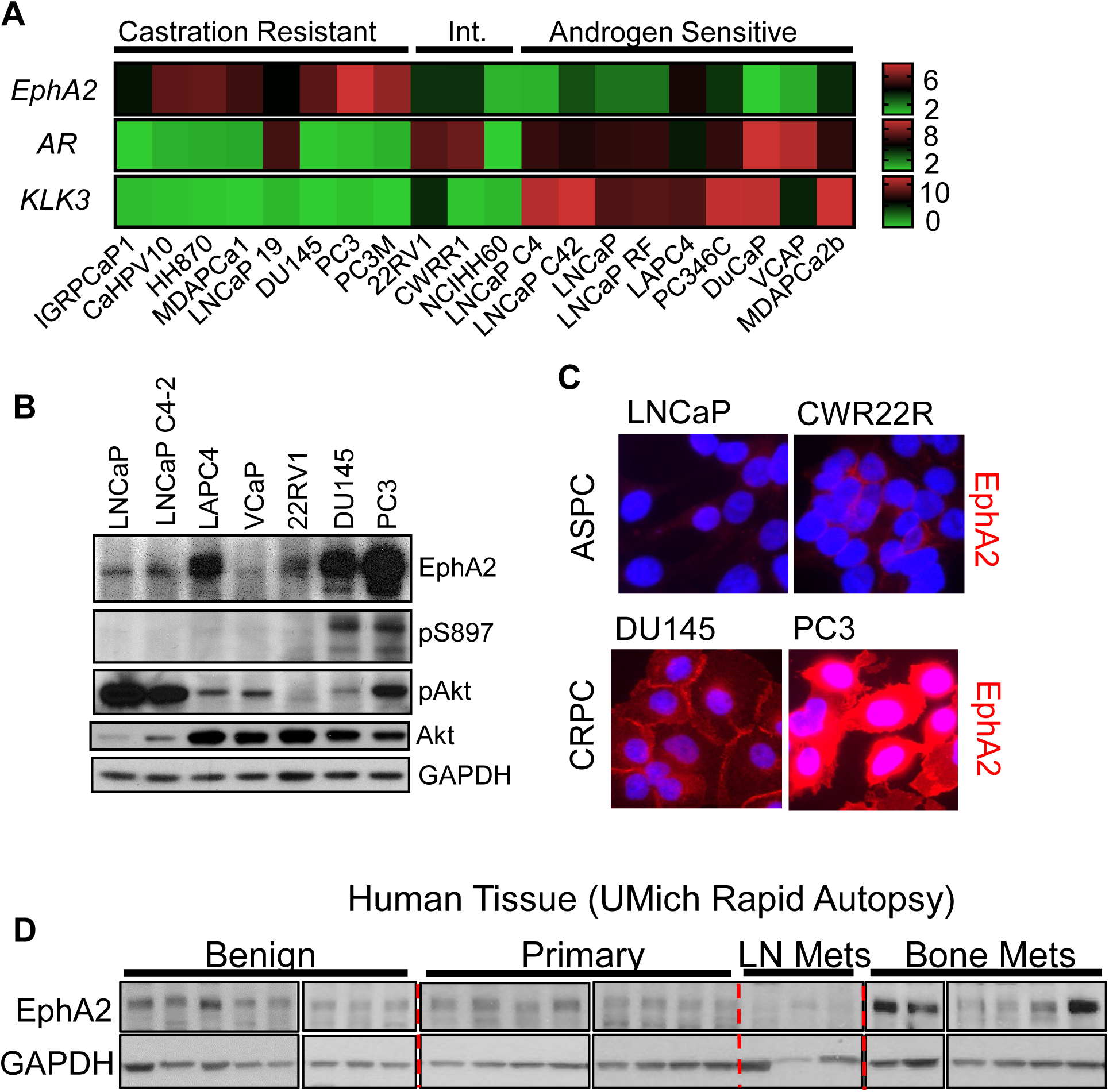
EphA2 is Overexpressed in Castration Resistant and Metastatic Prostate Cancer. **A)** Analysis of log_2_ transformed FPKM gene expression values extracted from bulk RNA seq published previously (Smith R. et al. *Scientific Reports* **2020**) shows high EphA2 expression in castration resistant, commercially available human PCa cell lines. **B)** Immunoblot characterization of select human PCa cell lines show increased EphA2 expression and S897 phosphorylation in mCRPC derived cell lines. **C)** EphA2 expression in representative human prostate cancer cell lines. **D)** Lysate of human prostate cancer tissues reveal increased expression of EphA2 in bone metastasis samples. Lysates were prepared from micro-dissected human prostate cancer tissues under the Rapid Autopsy Program at the Univ. Mich. PCa SPORE.

To further confirm EphA2 overexpression in late-stage human PCa tissues in pathologically relevant settings, we probed a panel of lysates prepared from microdissected primary and metastatic tissues harvested in the University of Michigan Rapid Necropsy Program. Interestingly, we found that EphA2 is more highly expressed in bone metastatic tissues compared to tissues from benign prostate, primary tumors, or lymph node metastases **(Fig. 1D)**.

Next, we investigated whether other members of the EphA receptors were similarly regulated in PCa. Surprisingly, among the 9 mammalian EphA receptors, EphA2 was the only one overexpressed in castration resistant cell lines in the Smith dataset (Rebecca Smith et al.) **(Fig. S1A)** indicating that EphA2 is the predominantly expressed EphA receptor in castration resistant PCa. Consistently, among all the mammalian EphA receptors, EphA2 has the strongest and most significant negative correlation to AR expression in human metastatic biopsies **(Fig. S1B)**.

### EphA2 is upregulated during malignant progression of PCa in vivo

To analyze the dynamics of EphA2 expression during PCa progression, we modeled increasing malignancy by serially passaging murine PCa cell lines in syngeneic host in vivo. We selected the TRAMP-C1 cell line derived from prostate adenocarcinoma induced by SV40 T antigen in C57Bl/6J mice^24,25^ Cells were injected subcutaneously; the resulting tumors were dissociated and cells were isolated for culture **(Fig. 2A)**. When re-grafted into new mice, the selected cells, termed in vivo 1 (IV1), exhibited accelerated tumor development compared to the parental TRAMP-C1 cells with five weeks of shortened latency **(Fig. 2B)**. Interestingly, the IV1 cells show significantly elevated expression of EphA2 **(Fig. 2C,D)**. The basal level of pS897 was also increased **(Fig. 2C,E)**, suggesting activation of pro-oncogenic noncanonical signaling by EphA2. The levels of pAkt and pERK, known upstream kinases that directly and indirectly phosphorylate S897, respectively, were also significantly increased **(Fig. 2C,F,G)**. In keeping with the inverse correlation between AR and EphA2 expression in human PCa cell lines, the more malignant TRAMP-C1 IV1 cells also showed loss of androgen receptor **(Fig. 2C,H)**. Here AR loss serves as a marker of progression toward CRPC coinciding with increased EphA2 expression, however, our investigation did not reveal a direct relationship between AR and EphA2 expression. Expression of E-cadherin, an epithelial cell marker highly expressed in parental cells **(Fig. 2C,I)**, is reduced to nearly undetectable level, consistent with increased malignancy of IV1 cells associated with epithelial-mesenchymal transition (EMT). The basal catalytic activation of EphA2 tyrosine kinase activity, as measured by pY588 did not change much **(Fig. 2C,J)** despite higher EphA2 expression, suggesting diminished canonical signaling.

**Fig. 2.**
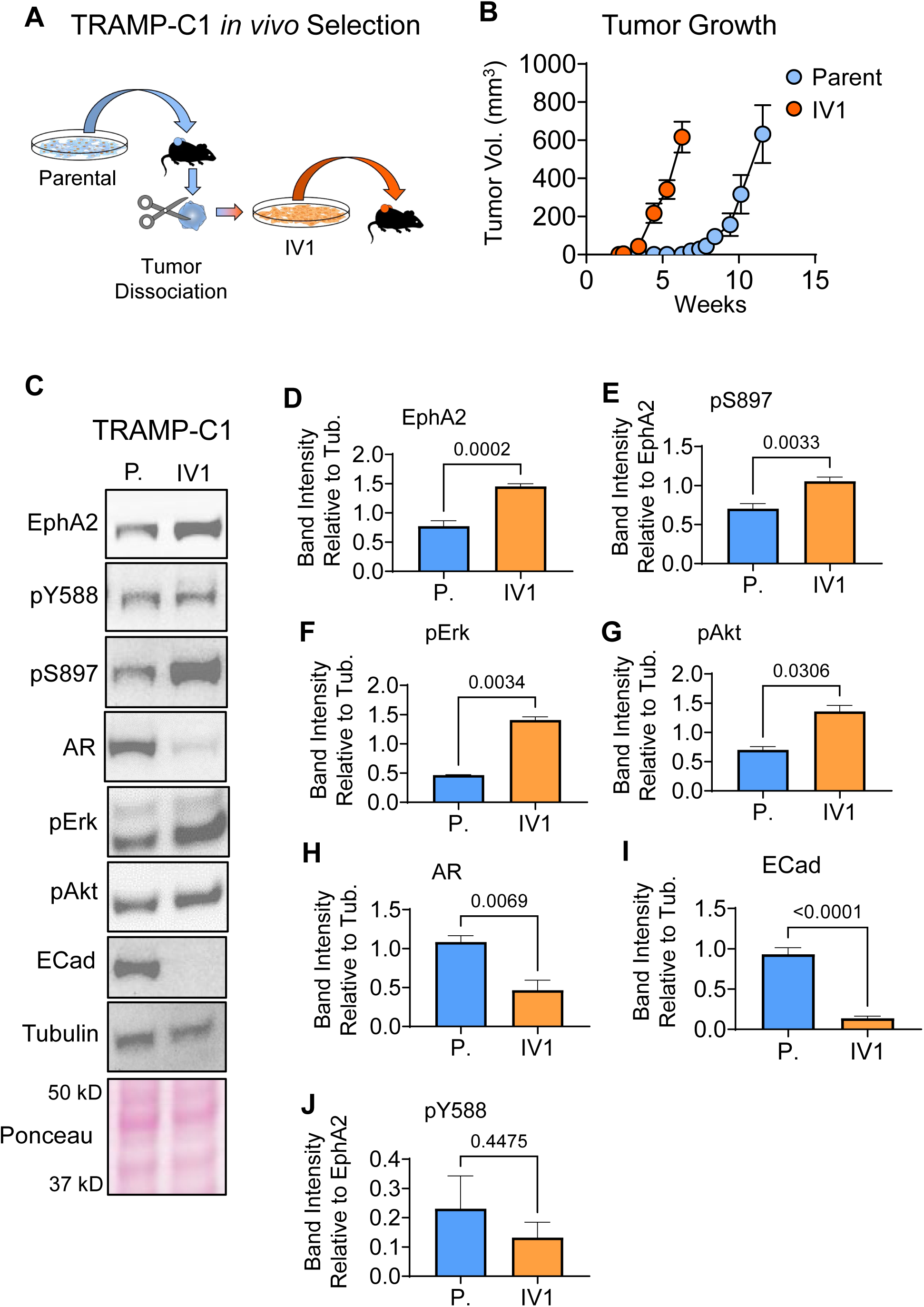
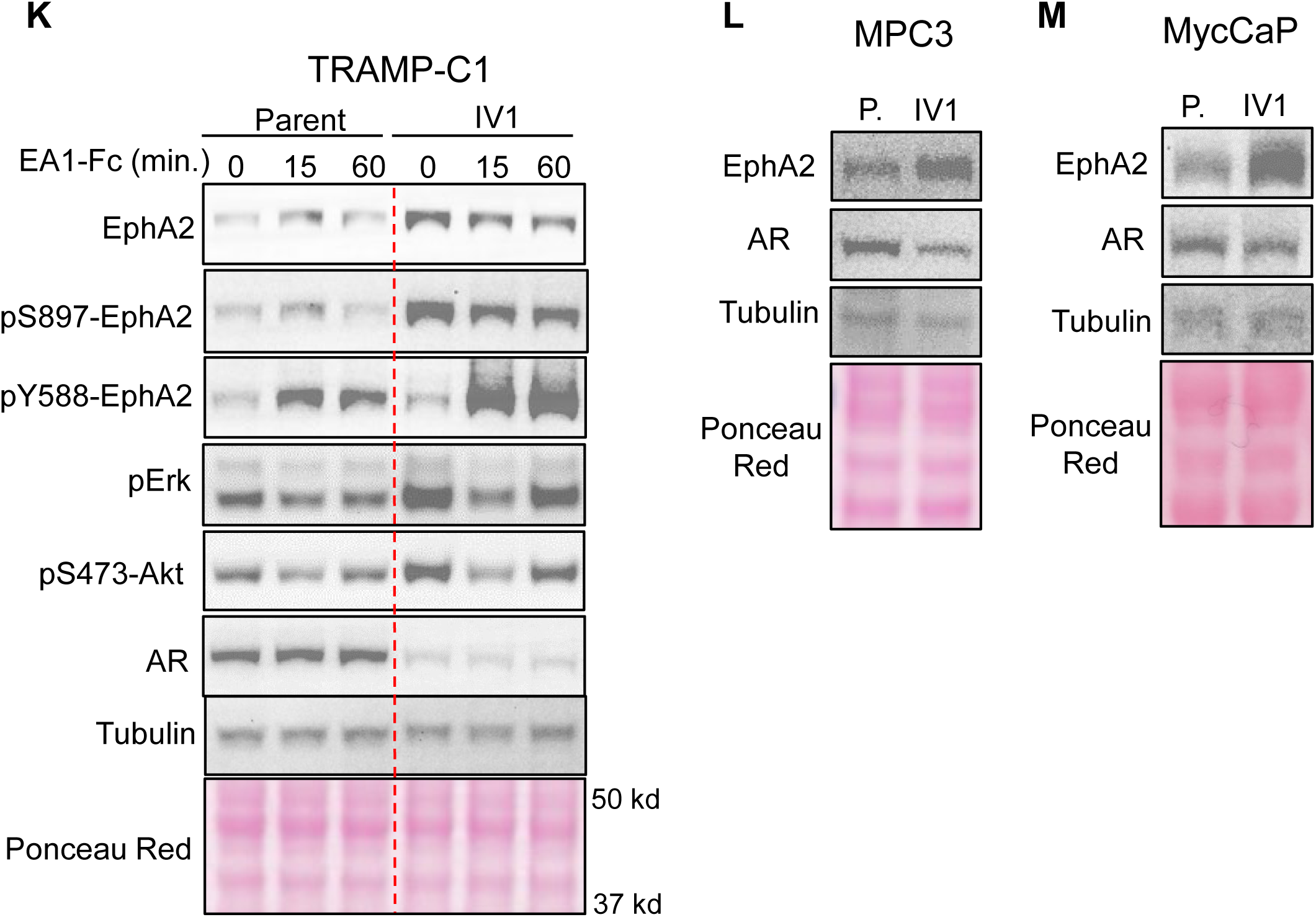
EphA2 is upregulated following in vivo selection for enhanced tumorigenicity. **A)** Diagram depicting the serial propagation of TRAMP-C1 cells in syngeneic mice to select for cells with increased tumorigenic capacity. **B)** The *in vivo* selection of cells resulted in accelerated tumor growth and reduced latency. **C)** Immunoblot analyses of the in vivo selected (IV1) and the parental TRAMP-C1 cells using the indicated antibodies. **D,E,F,G,H,I,J)** Quantification of relative band intensity (n = 4). All bands normalized to tubulin, except for pY588 and pS897 which are normalized to total EphA2 expression. **K)** Immunoblot of TRAMP-C1 parental and IV1 cells after stimulation with Ephrin-A1-Fc ligand for the indicated times. **L,M)** MPC3 and MycCaP cells were propagated in syngeneic C57Bl/6 and FVB mice, respectively. Lysates of the parental and the in vivo selected cells (IV1) were subject to immunoblot using the indicated antibodies.

As we have shown previously^10^, stimulation with exogenous ephrin-A1-Fc ligand caused tyrosine kinase activation, leading to inhibition of ERK and Akt activities **(Fig. 2K)**. S897 phosphorylation associated with noncanonical pro-oncogenic function of EphA2 was decreased by the ligand treatment. Therefore, the overexpressed EphA2 retains its intrinsic tumor suppressive function upon ligand stimulation.

To further validate the correlation between EphA2 expression and elevated pS897, pERK and pAkt, parental TRAMP-C1 cells were sorted into high and low EphA2 expressing populations. The high EphA2 expressing cells showed a higher basal level of S897, Akt and Erk phosphorylation, and were highly responsive to ligand stimulation **(Fig. S2A)**.

Next, we examined whether the upregulation of EphA2 following in vivo selection for malignant cells can be extrapolated to other syngeneic murine PCa models. MPC3 and MycCaP PCa cell lines are derived from prostate-targeted co-deletion of *PTEN* and *Trp53* and overexpression of Myc on C57Bl/6J and FVB background, respectively.^26,27^ They were subjected to the same selection procedures in syngeneic mice **(Fig. 2A)**. Similar to TRAMP-C1, MPC3 and MycCaP cells selected *in vivo* exhibited increased EphA2 expression **(Fig. 2L,M)**. There also marked reduction of AR in MPC3 cells, while only a milder reduction was observed in MycCaP cells, possibly due to the very high basal level AR expression in these cells. Moreover, MPC3 cells subjected to iterative rounds of *in vivo* selection exhibited progressive increase in EphA2 expression **(Fig. S2B)**. Taken together, EphA2 expression and oncogenic signaling are increased concomitant with the loss of AR expression following in vivo selection for more malignant phenotype.

### Bone metastases show preferential EphA2 overexpression and activation of noncanonical signaling

To investigate the pathological relevance of noncanonical signaling of EphA2 in human PCa, we obtained human prostate cancer tissue microarray (TMA) established by the Rapid Autopsy Program of the Pacific Northwestern Prostate Cancer Spore and stained them for pS897- and total EphA2. The TMA contains primary tumors as well as metastases to the lymph node and bone metastases, as well as soft tissue metastasis to the liver. Consistent with the biochemical analysis of tumor lysates from the Rapid Autopsy Program of University of Michigan PCa SPORE **(Fig. 1E)**, significantly higher EphA2 expression was detected in bone metastasis than the primary PCa or metastasis to the lymph and liver **(Fig. 3A, a-h** and **Fig. 3B)**. Furthermore, bone metastatic tumor samples exhibited significantly elevated EphA2-pS897 staining compared to primary PCa and metastases to the LN and liver **(Fig. 3A, i-p** and **Fig. 3C)**, demonstrating selective upregulation of pro-oncogenic, noncanonical signaling by EphA2 in bone metastasis relative to primary tumors and visceral metastases. Of note, we could not detect pS897 in the PCa tissue lysates used in **Fig. 1D**, possibly due to the highly labile nature of the phosphoserine signals that were lost during lysate preparation and/or storage. Tissues for TMA were immediately fixed after dissection and are more likely to retain pS897 signals. Confirming the biochemical **(Fig. 1D)** and IHC results **(Fig. 3A)**, analysis of the SU2C/PCF dataset revealed significantly higher EphA2 expression in bone metastases compared to all other metastatic sites **(Fig. 3D)**. More importantly, higher EphA2 expression is correlated with worse survival among patients with bone metastasis (**Fig. 3E**), whereas no such correlation was observed when all metastatic sites were included in the analysis **(Fig. S3A)**, nor was there such a correlation among primary PCa patients **(Fig. S3B)**.

**Fig. 3.**
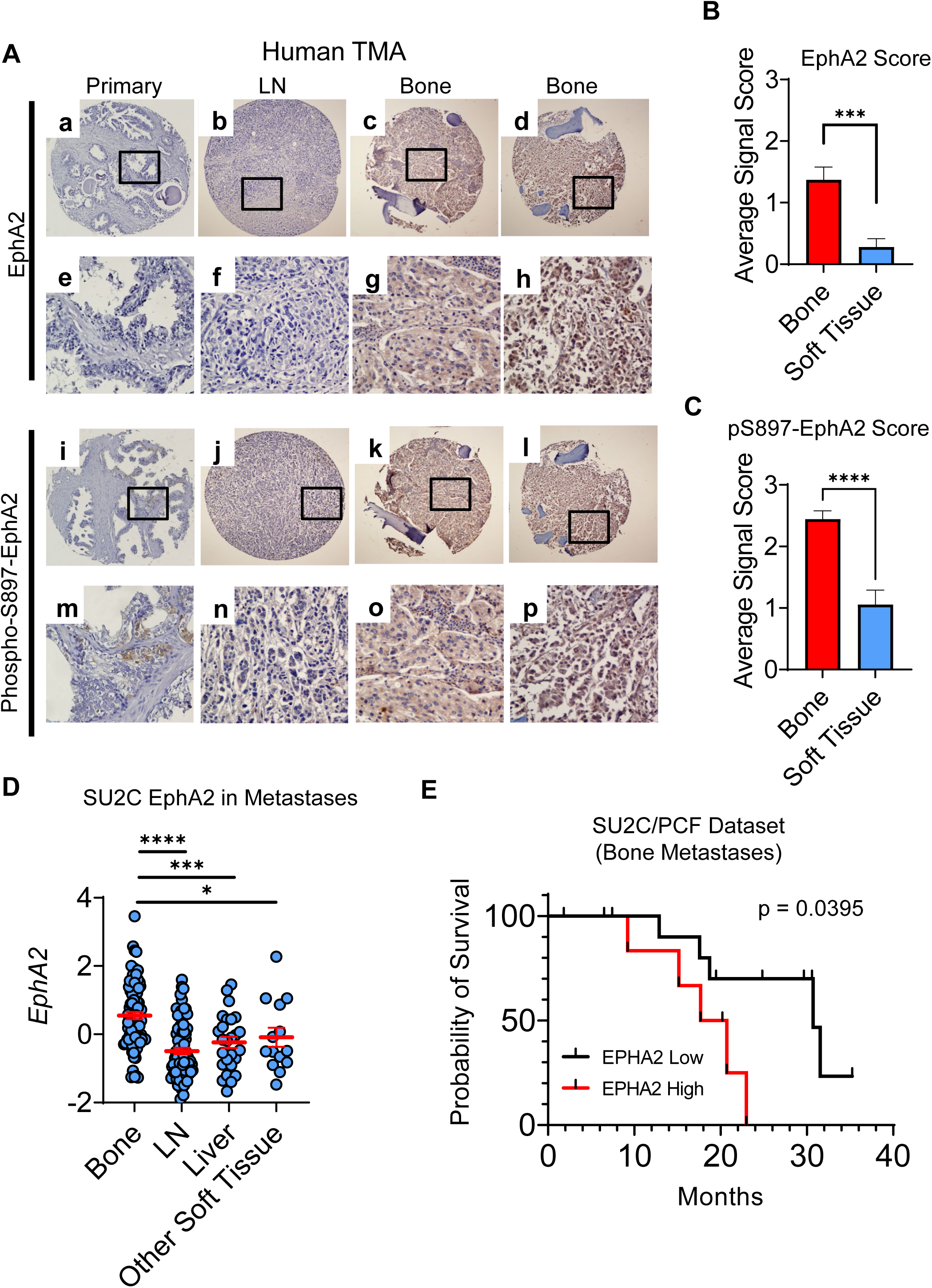
Differential overexpression and activation of noncanonical signaling of EphA2 in bone metastases. **A)** Immunohistochemical (IHC) analyses of tissue microarray (TMA) prepared from primary, lymph node metastatic (LN), and bone metastatic human PCa tissues from the Rapid Autopsy Program of the Pacific Northwest PCa SPORE spotted. **B and C)** The relative IHC staining strengths of EphA2 and pS897-EphA2 on TMA was scored using 0-3 scale (0, absent; 1, weak; 2, moderate; 3, strong) . Data consists of 27 bone metastatic samples, and 18 soft tissue samples including 10 lymph node, 5 liver, and 3 lung metastatic samples. Comparisons were made using an unpaired t-test (B, p = 0.0003 C, p < 0.0001) **D)** mRNA expression analysis of human PCa metastatic biopsies revealed that bone metastases have the highest EphA2 expression compared to other sites of disease dissemination. Data from 208 samples from the SU2C/PCF Dream Team. mRNA Expression z-scores relative to all samples (log FPKM Capture) accessed via cBioportal. **E)** Kaplan-Meier curve of patients with bone metastases according to the levels of EphA2 expression.

Most bone metastasis specimens came from patients with mCRPC. To test how acquisition of castration resistance impacts EphA2 noncanonical signaling, we used LAPC9 human prostate cancer model that is derived bone metastasis and can be propagated in vivo as xenograft in immune deficient mice. As shown in **Fig. S3C**,**D**,**E**, LAPC9 cells derived from the castrated host showed increased pS897 compared with androgen sensitive parental cells, as evidenced by both immunoblot and IHC staining. In sum, these data suggest that PCa metastases to the bone are associated with overexpression of EphA2 and activation of noncanonical signaling, which is correlated with worse prognosis of the patients.

### Disruption of noncanonical pro-oncogenic signaling via S897A mutation suppresses growth of PCa xenografts

To investigate the pathological relevance of EphA2 noncanonical signing through S897 phosphorylation in PCa, we expressed S897A mutant EphA2 that abolishes noncanonical signaling in PC-3 cells. WT EphA2 was used as control, and both were expressed at similar levels **(Fig. 4A)**. Upon serum stimulation of starved cells, we observed strong induction of pS897 on WT EphA2. S897A-EphA2 cells had pS897 level that was comparable to vector control cells, suggesting the weak signal likely originated from the endogenous EphA2. Upon implantation into nude mice, PC-3 cells expressing S897A-EphA2 mutant exhibited significantly diminished tumor development compared to either the WT-EphA2, or vector expressing tumors **(Fig. 4B)**.

**Fig. 4.**
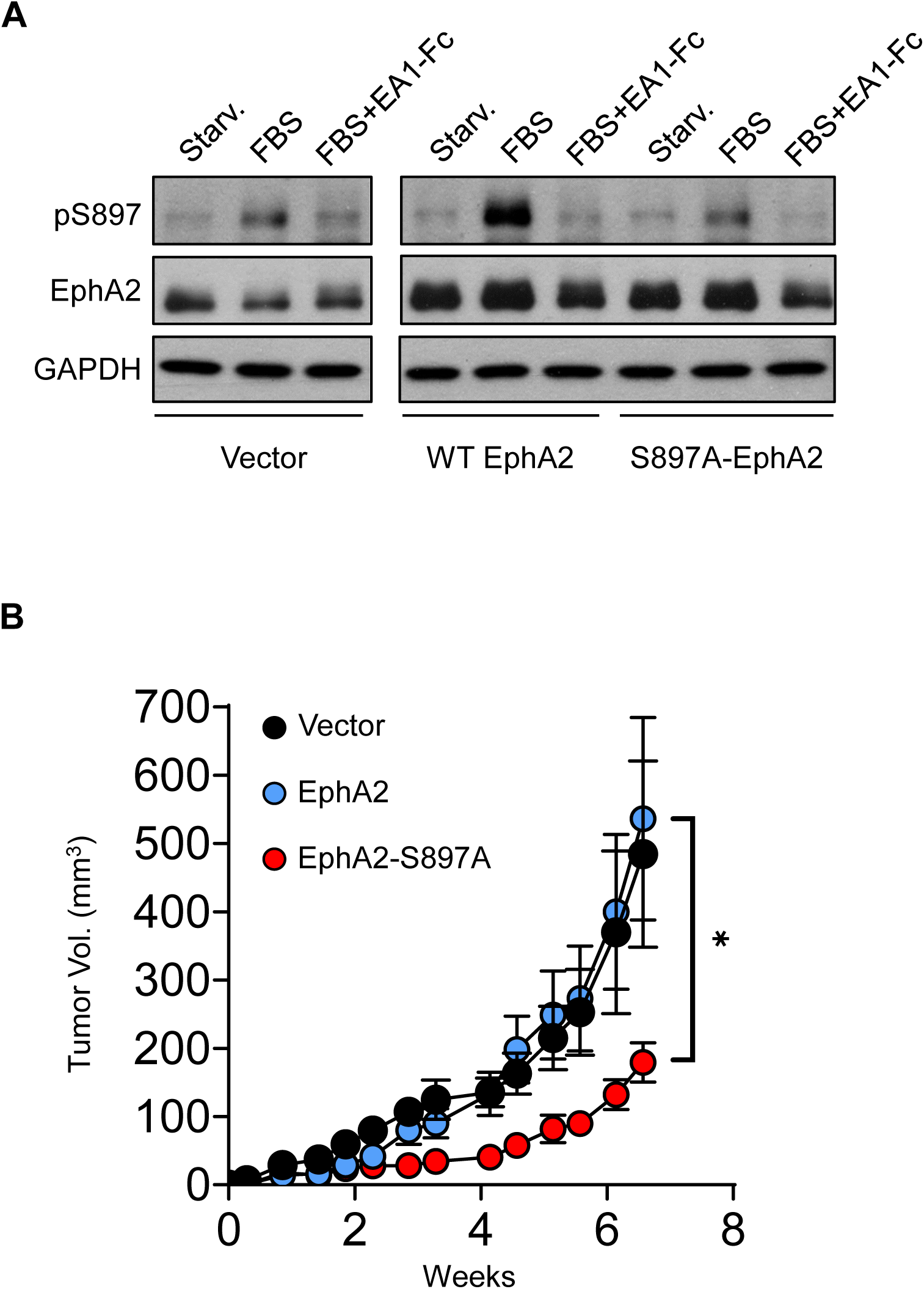
S897A mutation that disrupts the oncogenic noncanonical signaling resulted in diminished tumor development. **A)** Immunoblot analysis of PC3 cells transduced with lentivirus to express WT-EphA2 or S897A-EphA2. Empty vector-transduced cells were used as control. Cells were starved for 24 hours in serum-free medium and stimulated with either FBS alone or FBS togethr with ephrin-A1-Fc. **B)** PC3 cells overexpressing WT-EphA2 or S897A-EphA2 mutant were xenografted subcutaneously into nude mice (n = 10 per group). Vector alone cells were used as control. Comparison between WT-EphA2 and S897A-EphA2 made with an unpaired t-test at day 46 post-injection (p = 0.0402).

Next, we expressed WT or S897A-EphA2 in C4-2, an androgen insensitive human PCa cancer cell line derived from androgen-sensitive LNCaP. Like LNCaP, C4-2 cells express very low endogenous EphA2 **(Fig. 1A,B)**. Overexpression of WT EphA2 significantly promoted C4-2 xenograft tumor development, and the effect was diminished by S897A mutation **(Fig. S4A,B)**, indicating the requisite role of serine 897 phosphorylation in promoting tumorigenesis by EphA2.

### Lack of ligand expression in the bone provides a permissive milieu for PCa metastasis

PCa metastasis is known to have strong bone tropisms. In the castration-resistant setting, 70%–90% of patients have radiographically detectable skeletal involvement^28–30^. The selective upregulation of EphA2 expression and noncanonical signaling in skeletal metastases suggests a potential role in the bone tropism. One of the best characterized functions of Eph receptors is the repulsion of migrating cells upon contact with ligand-presenting cells. This prompted us to examine the expression of ephrin-A ligands in the bone microenvironment. Strikingly, analysis of the Human Protein Atlas shows undetectable or very low expression of all five mammalian *EFNA* genes in the bone, which contrasts with all other tissues examined that have high *EFNA1* expression and variable expression of the other four *EFNA* genes **(Fig. 5A-E)**. To further validate the observation, we obtained human bone marrows from four normal donors. qPCR analysis showed undetectable or very low expression of the five *EFNA* genes in all four bone marrow specimens relative to 22Rv1 human prostate cancer cell line with varying expression of *EFNAs* **(Fig. 5F)**. The absence of ephrin-As in the bone microenvironment creates a highly permissive microenvironment, as PCa cells presenting abundant EphA2 on the surface can home to the bone without being repulsed by the ligands on the skeletal cells. Moreover, the dearth of ephrin-A ligands in bone milieu together with their lack of expression on mCRPC cells (see below) also helps sustain the ligand-independent noncanonical signaling by EphA2 to promote PCa invasion and growth in the bone.

**Fig. 5.**
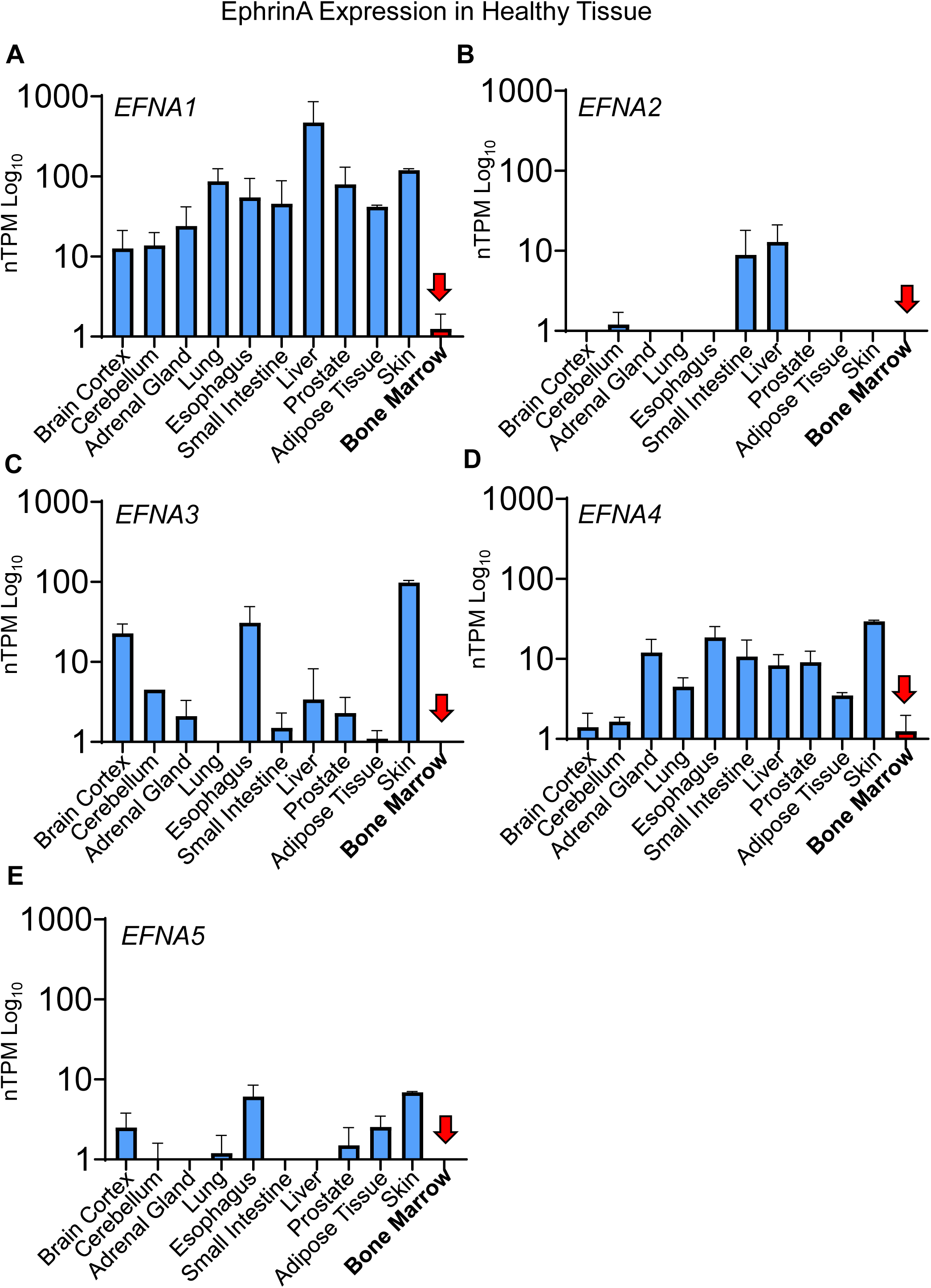
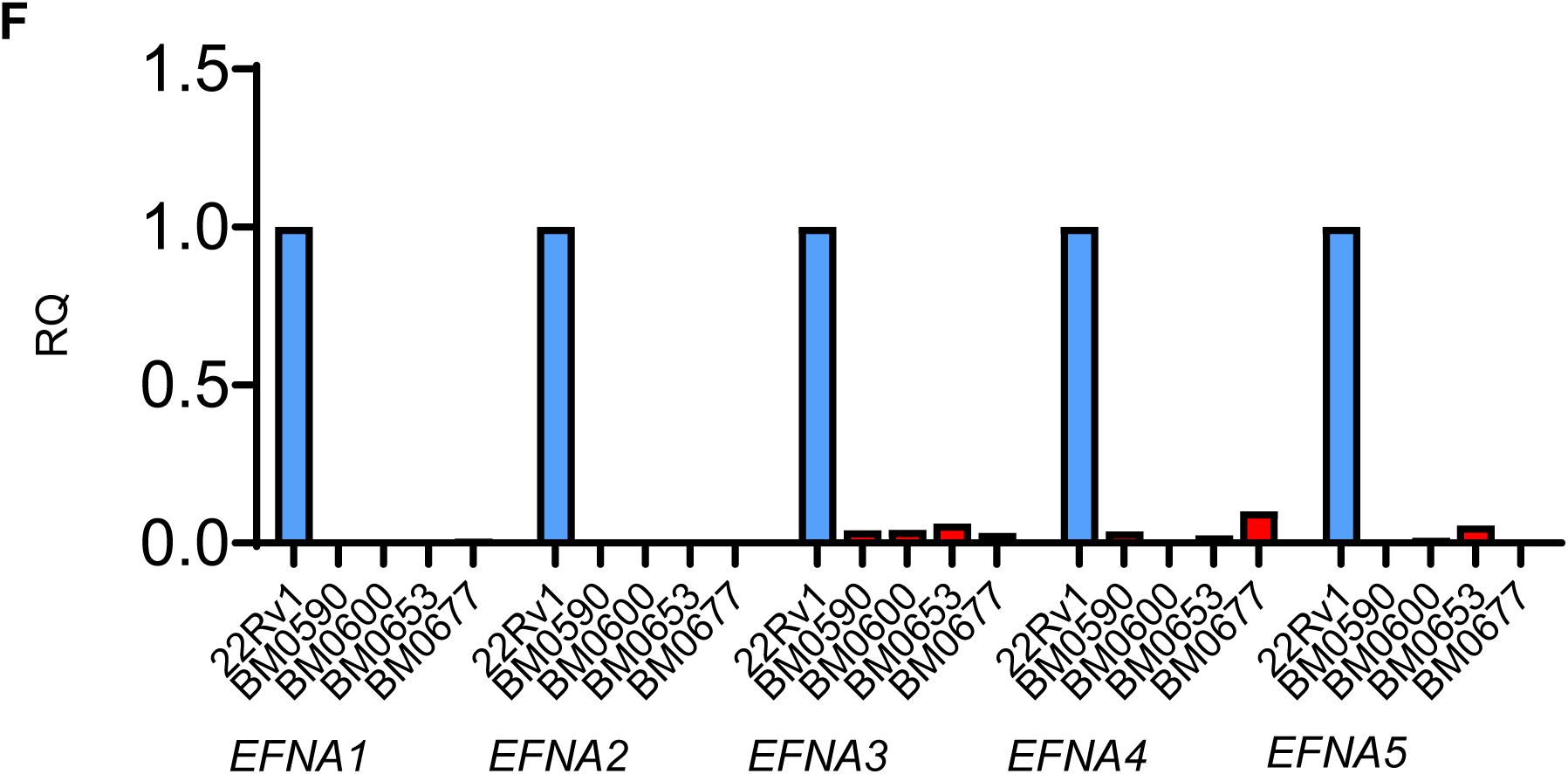
Undetectable or very low expression of all five *EFNA* genes in the bone microenvironment. **A-E)** EFNA1,2,3,4,5 expression across human tissues. Data was collected from Human Protein Atlas. **F)** qPCR analysis of bone marrow samples from four healthy human donors; 22Rv1 cells were used as control.

### Ephrin-A1 expression is lost in mCRPC

The duality of EphA2 as a tumor suppressor gene and as an oncogene is dictated by the presence and absence of the ephrin-A ligands, respectively. While the lack of the ephrin-A ligands in the bone marrow provides a permissive milieu, the ephrin-A ligands expressed on tumor cells can interact with EphA2 on adjacent cancer or stromal cells to trigger receptor-ligand interactions *in trans* and initiate tumor suppressive canonical signaling. Conversely, loss of ligand can facilitate pro-oncogenic EphA2 noncanonical signaling. Previous studies using mCRPC from rapid tissue acquisition necropsy show that *EFNA1* is one of three genes whose expression was significantly reduced in bone metastases compared with primary tumors and lymph node or liver metastasis.^17^ Consistently, we observe a low mRNA expression of all five *EFNA* genes across multiple castration resistant human PCa cell lines compared to androgen sensitive cells **(Fig. 6A)**. *EFNA1* gene is widely expressed in all androgen sensitive cell lines followed by *EFNA3,4,5*. Immunoblot results confirmed the loss of ephrin-A1 expression in PC-3 and DU-145 CRPC cells compared with androgen sensitive cell lines **(Fig. 6B,C)**. Moreover, analysis of the MSK database (*Cancer Cell*, **2010**) showed that low *EFNA1* expression is associated with worse patient survival **(Fig. 6D)**. These EphA2 overexpressing and ephrin-A deficient prostate cancer cells can be preferentially selected in the bone as described above.

**Fig. 6.**
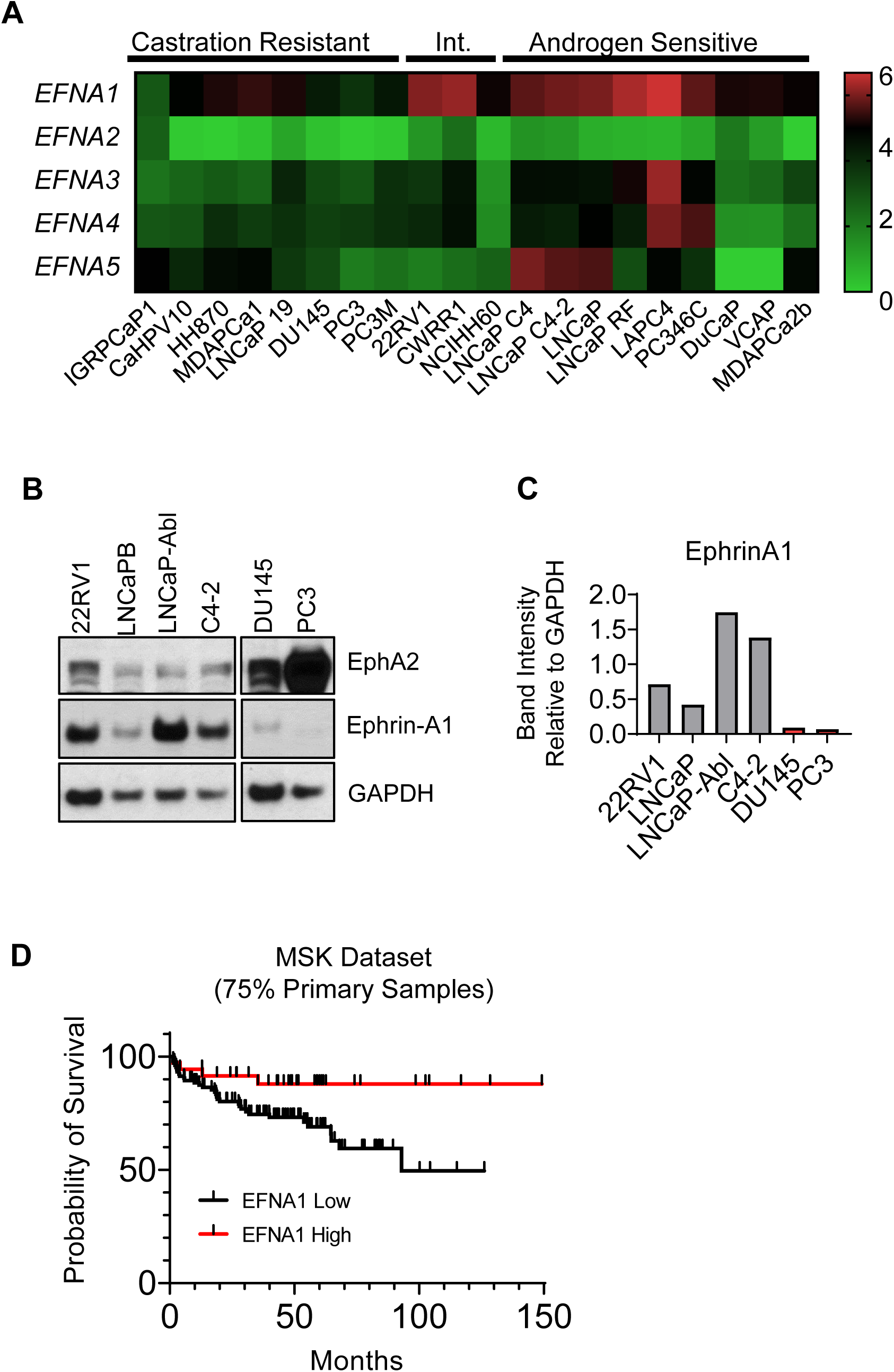
Ephrin-A1 is lost in mCRPC. **A)** Analysis of log_2_ transformed FPKM gene expression values extracted from bulk RNA seq published previously (Smith R. et al. *Scientific Reports* **2020**) showing low *Efna1-5* gene expression in commercially available castration resistant and androgen sensitive human PCa cell lines. **B)** Immunoblot characterization of select PCa cell lines reveals simultaneous high EphA2 expression and low EphrinA1 expression in metastatic castration resistant cells compared with androgen sensitive cells. **C**) Quantitative analysis of data in (B). **D)** Kaplan-Meier survival curve from the MSK dataset (*Cancer Cell* **2010**). Samples were aggregated into the top 25 percent EFNA1 expressing (*EFNA1* High), and lower 75 percent *EFNA1* expressing (*EFNA1* Low). Data were accessed via cBioportal.

### Restoration of *EFNA1* expression in PC-3 cells suppresses oncogenic noncanonical signaling by EphA2 and reduces tumor development

The results above suggest that the loss of ephrin-A1 expression and ensuing activation of non-canonical signaling by the overexpressed EphA2 may contribute to malignant progression of PCa. To directly test this hypothesis, we used a lentiviral vector to re-introduce *EFNA1* expression into PC-3 cells that are originally derived from a bone metastasis and have little expression of the five *EFNA* genes. As shown in **Fig. 7A**, restoration of ephrin-A1 resulted in dramatic reduction in the total level of EphA2, likely due to EphA2 activation by ligand presented *in trans* on adjacent cells, leading to endocytosis and proteolytic degradation of the receptor. Ephrin-A1 re-expression also led to proportionally stronger basal tyrosine phosphorylation in the juxtamembrane domain (pY588) and reduced pS897 **(Fig. 6B,C)**, which is associated with canonical tumor suppressive, and noncanonical oncogenic signaling of EphA2, respectively.

**Fig. 7.**
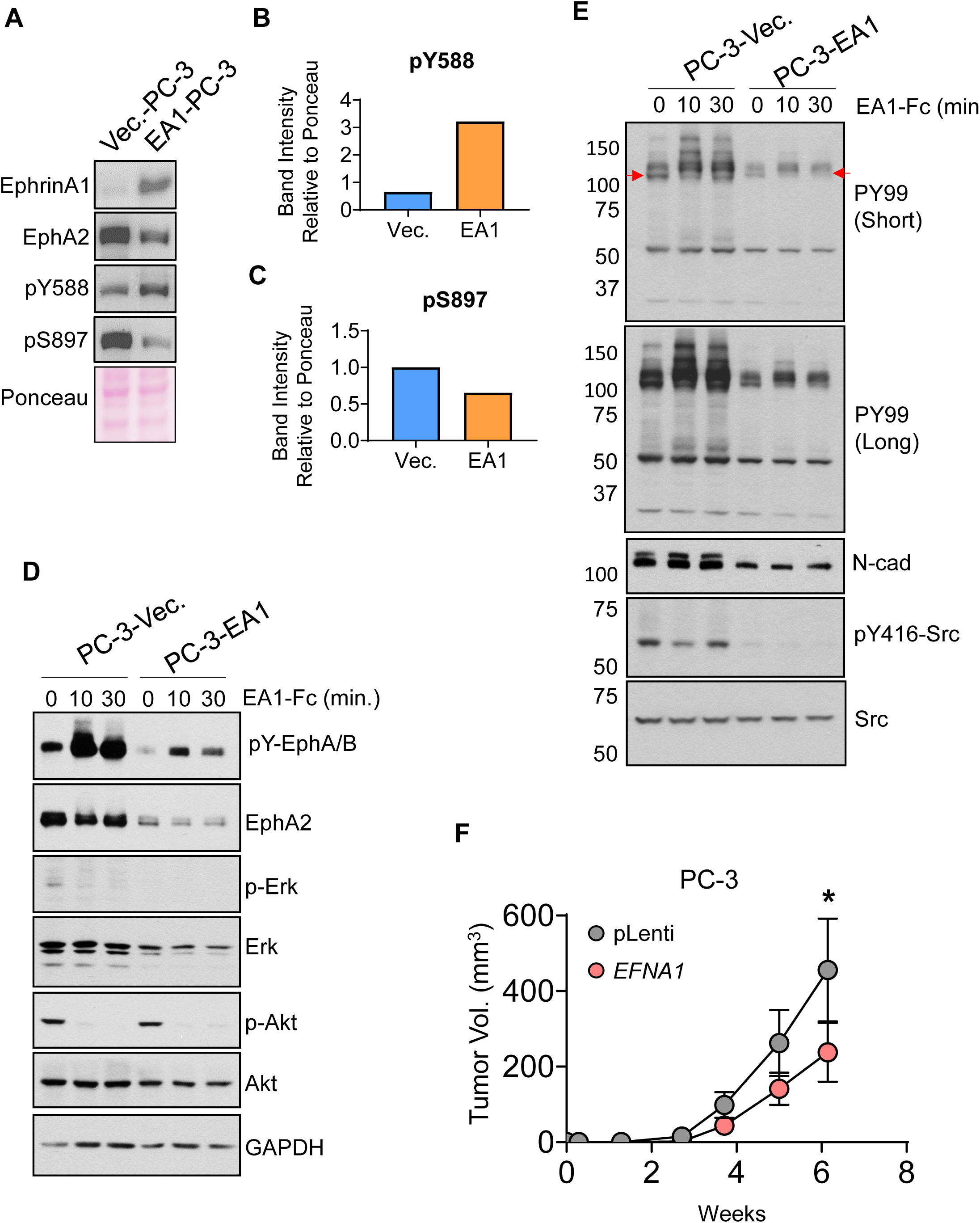
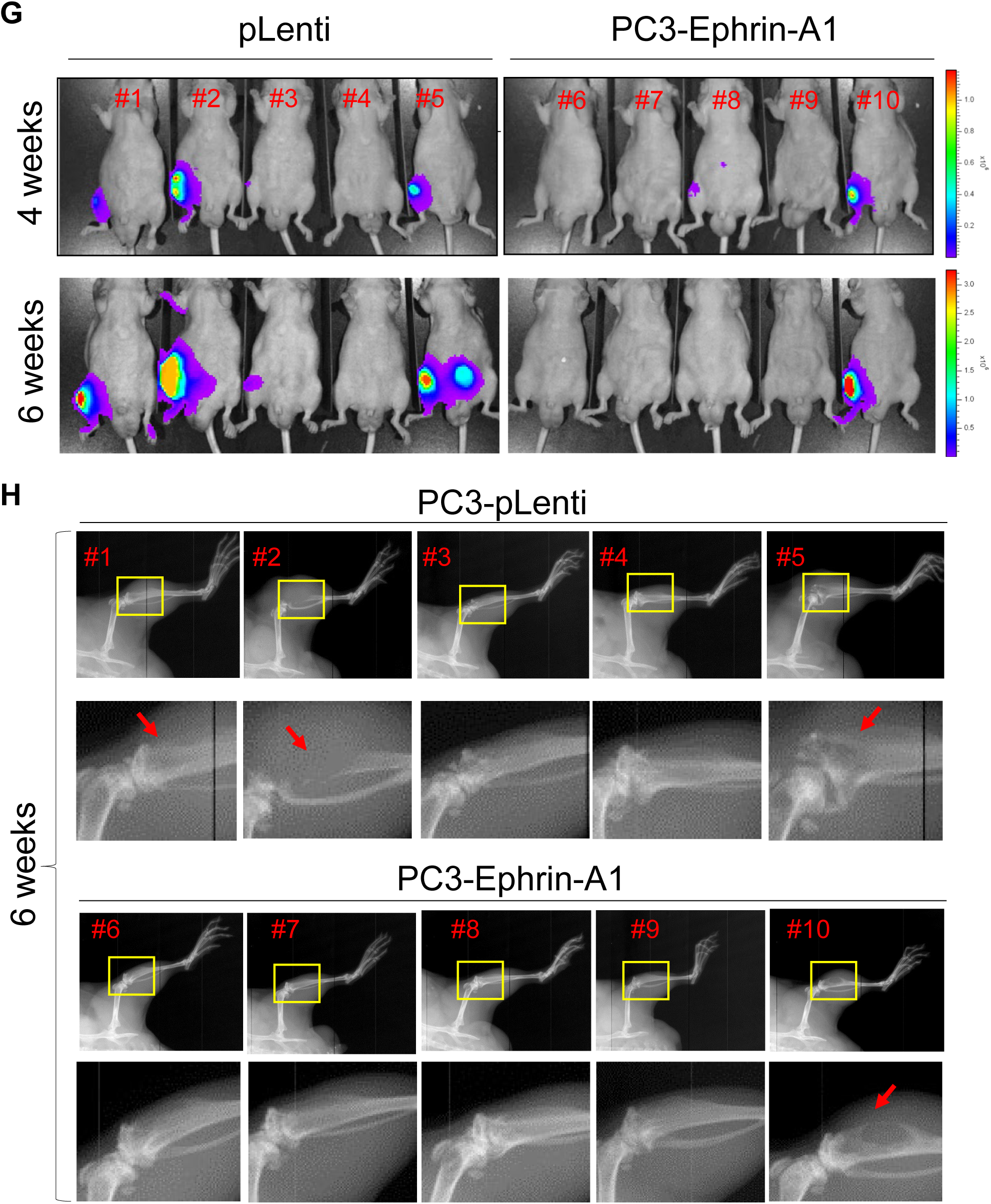
Restoration of *EFNA1* expression in PC-3 cells suppressed oncogenic noncanonical signaling by EphA2 and inhibited subcutaneous and skeletal tumor development. **A)** Lysates of PC-3 cells transduced by lentiviral vector expressing *EFNA1* (EA1) or empty control vector were subject to immunoblot with the indicated antibodies. **B,C)** Quantification of pY588 marking canonical signaling and pS897 marking noncanonical signaling by EphA2, relative to total EphA2 expression. **D)** Vec. and EA1-expressing PC-3 cells were stimulated with recombinant exogenous EA1-Fc ligand *in vitro* for the indicated times, and lysates were subject to immunoblot with the indicated antibodies. **E)** PY99, a monoclonal antibody that recognizes most pY motifs, was used to probe overall pY profiles using lysates from Vec. and EA1-expressing PC-3 cells stimulated with EA1-Fc. N-cadherin and pY416-Src were also probed. **F)** PC-3 cells expressing EA1 exhibit reduced cell growth in an in vitro cell proliferation assay. **G)** Vec. control and EA1-expressing cells were implanted subcutaneously into nude mice. Paired t-test to be used to monitor (p = 0.0319). **G)** Bioluminescence imaging of mice four and six weeks after intratibial injection of PC3-vec. or PC-3-EA1 expressing cells tagged with luciferase. **H)** X-ray imaging of mice 6 weeks post intratibial injection of PC3-Vec and PC3-EA1 cells.

As reported previously,^10,11^ stimulation of control cells with recombinant ephrin-A1-Fc led to potent activation of canonical signaling characterized by EphA2 tyrosine phosphorylation and suppression of both ERK and Akt **(Fig. 7D)**. Restoration of ephrin-A1 in PC-3 cells led to diminished basal ERK activities **(Fig. 7D**, lanes 1 vs. 4). The basal activities of Akt, on the other hand, remained unchanged. Despite much lower basal level of EphA2 on ephirn-A1-restored cells, it remained responsive to exogenous ligand stimulation, leading to inhibition of Akt **(Fig. 7D)** and pS897 **(Fig. S5)**.

To explore how ephrin-A1 restoration effect global changes in cell signaling machinery, we probed whole cell lysates for overall pY profile utilizing an anti-pY antibody (PY99) that recognized most tyrosine-phosphorylated proteins. Stable restoration of ephrin-A1 caused a profound reduction in total pY involving many proteins across a large molecule weight range **(Fig. 7E**, lanes 1 vs. 4, long exposure), demonstrating the powerful roles of ephrin-A1-induced activation of EphA2 in prostate cancer cell regulation. While the identities and functions of the tyrosine phosphorylated proteins remained to be characterized, Src was one of the molecules affected, exhibiting a robust reduction in pY416-Src **(Fig. 7E**, lanes 1 vs. 4), the active form of the kinase.

Stimulation with exogenous recombinant ephrin-A1-Fc led to massive upregulation of pY in vector control cells **(Fig. 7D)**. Interestingly, ligand treatment also caused tyrosine de-phosphorylation of some proteins **(Fig. 7E**, red arrow), including transient dephosphorylation of Src **(Fig. 7E**, lane 1 vs. 2). The results indicate dynamic nature of EphA2 canonical signaling in controlling pY.

Expression of N-cadherin, a marker of epithelial-mesenchymal transition or EMT, was reduced in ephrin-A1-restored PC-3 cells, suggesting reversion to a less malignant phenotype. To test the effects *in vivo*, we implanted the vector and *EFNA1*-expressing PC-3 cells subcutaneously into immune-deficient nude mice. We found that ephirn-A1-PC-3 cells exhibited significantly diminished tumor growth relative to the vector control **(Fig.7G)**. In aggregate, these data demonstrate the critical importance of ephrin-A1 expression and the loss of it during tumor progression in promoting malignant behaviors of prostate cancer cells.

Next, we examined whether *EFNA1* ligand restoration could impact tumor development in the bone. Control and ephrin-A1-restored PC-3 cells were tagged with luciferase, and injected to the tibia of nude mice. In the control group, three of the five mice showed bioluminescent imaging (BLI) signals by 4 weeks **(Fig. 7G)**. The tumors continued to grow; by sixth week, the BLI signals grew stronger. Ephrin-A1 restoration suppressed bone xenograft development, and only one site showed BLI signal at six weeks. One lesion at week four (#8) disappeared by week 6 **(Fig. 7G)**. X-ray imaging of the tibia showed evidence of bone loss (red arrows) in three mice in the control group, while one mouse displayed moderate bone loss **(Fig. 7H)**. The intratibial injection of PC-3 was an imperfect model for bone metastases, as it did not happen spontaneously. However, the data did point to the importance of ligand loss on cancer cells in bone colonization.

In aggregate, the comprehensive characterization of EphA2-ephrin-A system in PCa using in vitro, in vivo model systems, as well as human patient specimens from two independent Rapid Autopsy Programs demonstrates important roles of the ligand loss-enabled activation of EphA2 noncanonical signaling via S897 in promoting PCa development. The overexpression of EphA2 and loss of ligand in mCRPC offers opportunities for development of novel therapeutic strategies targeting this receptor and ligand pair.

## Discussion

EphA2 has come under intense investigation due to its overexpression at the late stages of many human solid tumors, including prostate cancer. However, the pathological processes it regulates, and the underlying molecular and cellular mechanisms remain incompletely understood. In this study, we show that EphA2 is selectively overexpressed in metastasis to the bone, but not visceral metastasis. Serine 897 phosphorylation that mediates the ligand-independent noncanonical signaling by EphA2 was also upregulated in bone metastasis. Loss of ephrin-A ligands on cancer cells and tumor microenvironment promotes noncanonical signaling. We find that the bone is unique among many tissues examined in expressing little of the five *EFNA* genes, providing a permissive microenvironment for the disseminating cancer cells to colonize. Consistently, analysis of human datasets shows EphA2 overexpression is associated with skeletal but not visceral metastasis, whereas loss of ephrin-A1 expression in primary PCa is correlated with poor prognosis. Further, S897A mutation that ablates EphA2 noncanonical signaling, suppressed PCa development. Restoration of ephrin-A1 expression in PC-3 cells, originally derived from bone metastasis and devoid of ephrin-As, suppressed tumor development in vivo. Together these results suggest that overexpression of EphA2 and concomitant loss of ligands in PCa lead to activation of noncanonical signaling, while the dearth of ephrin-A ligands in the bone creates a permissive microenvironment to augment skeletal metastasis.

Metastatic prostate cancer is a leading cause of cancer-related morbidity and mortality in men. Clinically, it is often manifested following androgen deprivation therapy (ADT), which leads to metastatic castration-resistant prostate cancer (mCRPC). Metastasis of prostate cancer differentially targets the bone, including vertebrae that causes pathologic fracture and spinal cord compression leading to significant neurologic and functional disability. Meta-analysis of human prostate cancer show that around 73% mCRPC patients have bone with or without LN metastases, followed by visceral disease (20.8%) and LN-only disease (6.4%).^31^ While visceral metastasis has been associated with worse overall survival, the prevalence of skeletal disease and its hitherto incurable nature make it a major health burden in men. It is widely believed that bone microenvironment lends support for mCRPC, as a result of complex interactions between the microenvironment and tumor cells. However, the exact molecular and cellular nature of such interactions remains elusive. We find that the bone is quite unique in expressing little of the five ephrin-A ligands in mammalian system, while widely expressed all other tissues examined. The lack of ephrin-As in the bone has several implications in the context of colonization and proliferation of disseminating prostate cancer cells. First, since ephrin-A ligands are anchored to the cytoplasmic membrane by GPI, EphA-ephrin-A interactions mediate cell-cell contact signaling. One of the best characterized functional outcomes of the EphA-ephrin-A interactions is the mutual repulsion between the ligand- and receptor-presenting cells. Therefore, the dearth of ephrin-A ligand in the bone will present little resistance to the incoming prostate cancer cells overexpressing EphA2. Second, as we reported previously, ligand-induced activation of canonical signaling of EphA2 leads to strong inhibition of ERK and Akt activities, leading to reduced cell migration and proliferation.^11,12^ Without the ligands to trigger tumor suppressive canonical signaling, bone microenvironment can facilitate growth and colonization of disseminated prostate cancer cells. Finally, prostate cancer progression is associated with overexpression of EphA2 and concomitant loss of ligands, conditions known to activate the ligand-independent noncanonical EphA2 signaling that enhances cancer stem cell properties and drives invasion and proliferation.^9,10,32,33^ We propose that cancer cell-intrinsic changes in the balance between EphA2 and ephrin-A expression propels the selective homing and colonization of these cells in the bone. Consistent with this notion, liver expresses high levels of multiple ephrin-As and therefore represents a hostile environment for EphA2-overexpssing cells. Indeed, examination of hepatic metastases of prostate cancer collected from two Rapid Autopsy Programs showed little EphA2 expression, suggesting that the visceral disease has adopted alternatively mechanisms to take hold in the liver.

In vitro investigations reveal potential mechanisms on how the loss of ligand expression impacts the behavior of prostate cancer cells. Thus, in keeping with our previous studies,^11,12^ restoration of ephrin-A1 expression in metastatic PC-3 cells induced ligand-dependent canonical signaling characterized by: 1) activation of the tyrosine kinase function of EphA2 marked by pY588, concomitant with degradation of the receptor, a common mechanism of desensitization for most RTKs, 2) suppression of the oncogenic noncanonical signaling evidenced by reduced pS897, 3) inhibition of basal activation of ERK, but not Akt, and 4) reduction of N-cadherin expression, a hallmark of epithelial-mesenchymal transition. The four outputs are all in keeping with the tumor suppressor roles of EphA2 upon ligand-induced canonical signaling. Strikingly, we also observed profound changes in tyrosine phosphorylation (pY) of many proteins. While the broad trend is massive reduction of pY of many proteins, there was also notable reduction of pY in other proteins, including Src upon ligand stimulation. While the identity and function for many of these proteins in relation to EphA2 expression and signaling remain to be further investigated, the results highlight dramatic effects of ephrin-A ligand expression, or the loss of it, on global signaling and function of prostate cancer cells.

In vivo, ephrin-A1 restoration in PC-3 cells significantly impaired tumor development both subcutaneously and after implantation in the tibia. The latter was an imperfect model for bone metastases, as it did not happen spontaneously. However, the data did point to the importance of ligand loss on cancer cells in bone colonization. Supporting this notion, analysis of prostate cancer datasets from Memorial Sloan Kettering Cancer Institute showed significantly worse survival in patients who have lost ligand expression in primary prostate cancer.

Xenograft studies also demonstrated the importance of pS897 in promoting tumor development by the overexpressed EphA2, as S897A mutation that blocks S897 phosphorylation reduced growth of both PC-3 and C4-2 xenografts. Staining of metastasis tissue array prepared from patient specimens of the Rapid Autopsy Program showed strong pS897 signals. Together these results indicate the pathological significance of the noncanonical signaling of EphA2 via pS897 in driving metastatic progression of prostate cancer.

Our findings highlight the translational significance of EphA2 in skeletal metastases of prostate cancer. As a type I transmembrane protein overexpressed on cell surface, EphA2 is amenable to a variety of targeting strategies. One strategy is the development of small molecules or monoclonal antibodies that induced the catalytic activation of EphA2, which is expected to restore its intrinsic tumor suppressor function. Indeed, we have previously discovered Doxazosin, an inhibitor of α1 adrenergic receptor still used in the clinic, is a novel agonist for EphA2 that was able to suppress PC-3 xenograft metastasis after orthotopic injection.^34,35^ Other approaches include induced degradation to the receptor. Future development of the novel EphA2-targeted agents could provide new drugs to tame the largely incurable skeletal disease of prostate cancer.

