## Supplementgal Figures for "Skeletal Metastasis of Prostate Cancer Is Augmented by Activation of EphA2 Noncanonical Signaling and Ligand-Deficient Bone Microenvironment"

### Fig. S1.

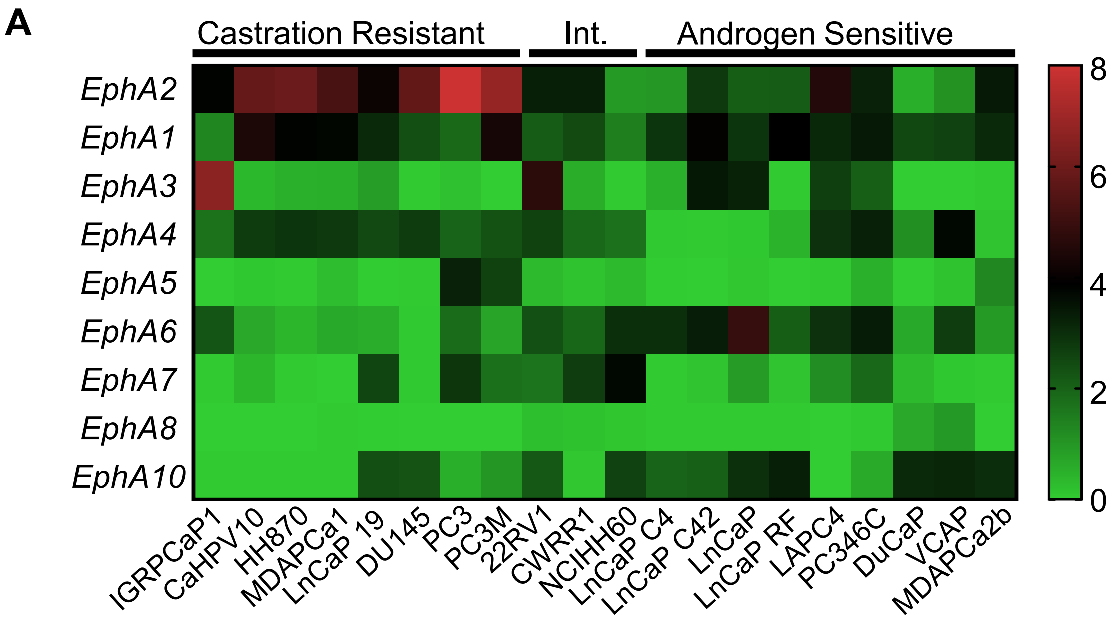
A
Castration Resistant
Int.
Androgen Sensitive
EphA2
EphA1
EphA3
EphA4
EphA5
EphA6
EphA7
EphA8
EphA10
IGRPCaP1
CaHPV10
HH870
MDAPCa1
LnCaP 19
DU145
PC3
PC3M
22RV1
CWRR1
NCIHH60
LnCaP C4
LnCaP C42
LnCaP
LnCaP RF
LAPC4
PC346C
DuCaP
VCAP
MDAPCa2b
8
6
4
2
0

**Fig. S1.**

**B**

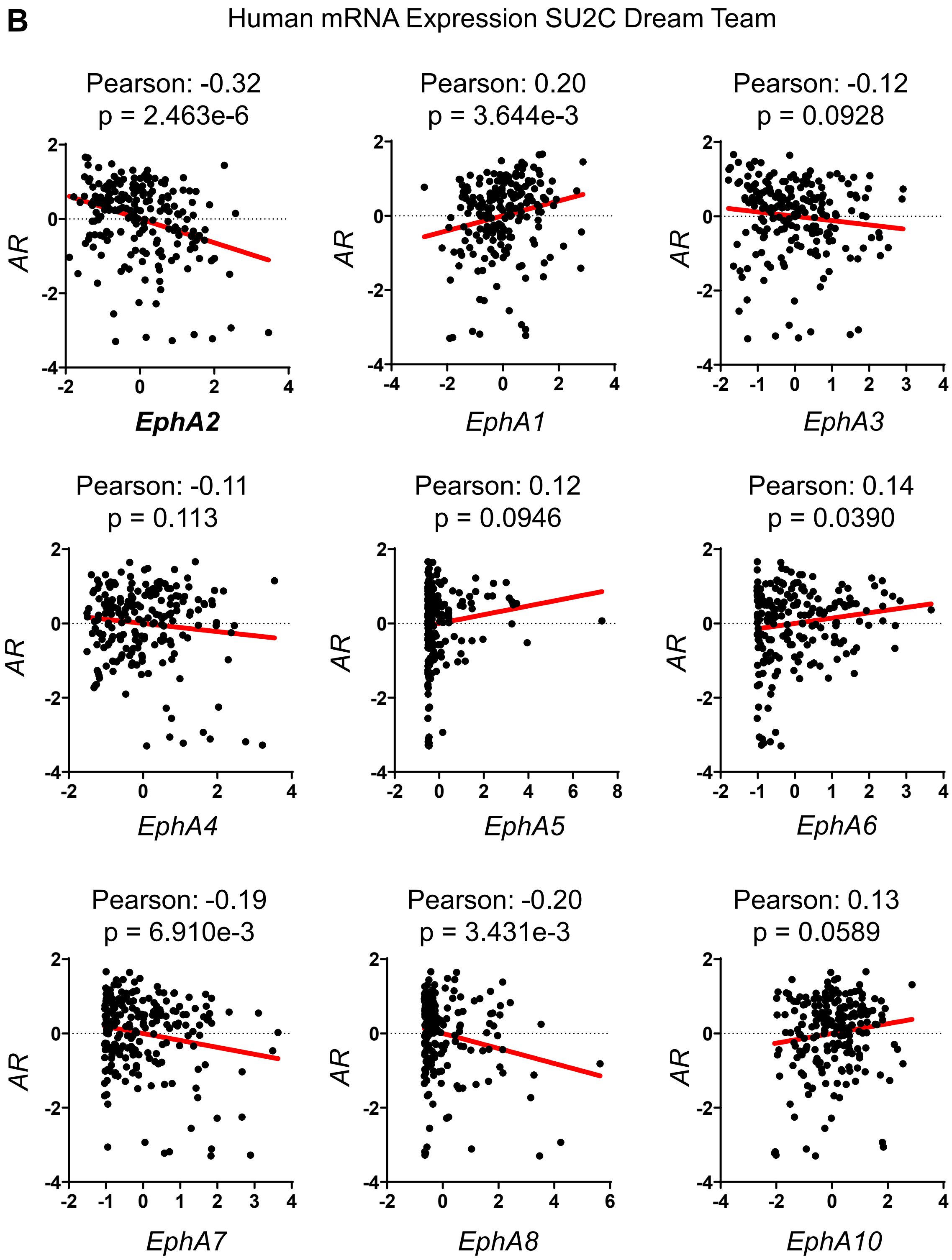
Human mRNA Expression SU2C Dream Team
Pearson: -0.32
p = 2.463e-6
EphA2
Pearson: 0.20
p = 3.644e-3
EphA1
Pearson: -0.12
p = 0.0928
EphA3
Pearson: -0.11
p = 0.113
EphA4
Pearson: 0.12
p = 0.0946
EphA5
Pearson: 0.14
p = 0.0390
EphA6
Pearson: -0.19
p = 6.910e-3
EphA7
Pearson: -0.20
p = 3.431e-3
EphA8
Pearson: 0.13
p = 0.0589
EphA10
AR

**Fig. S1. EphA2 is the Predominantly Expressed EphA Receptor in mCRPC. A)** Analysis of $log_2$ transformed FPKM gene expression values extracted from bulk RNA seq published previously (Smith R. et al. *Scientific Reports* **2020**) shows that EphA2 is the predominately expressed EphA receptor in human castration resistant PCa cell lines. **B)** EphA2 has the strongest and most significant negative correlation to AR expression of all the EphA receptors in metastatic human biopsies based on data is from the SU2C/PCF Dream Team. mRNA Expression z-scores relative to all samples (log FPKM Capture) accessed via cBioportal.

**Fig. S2.**

A

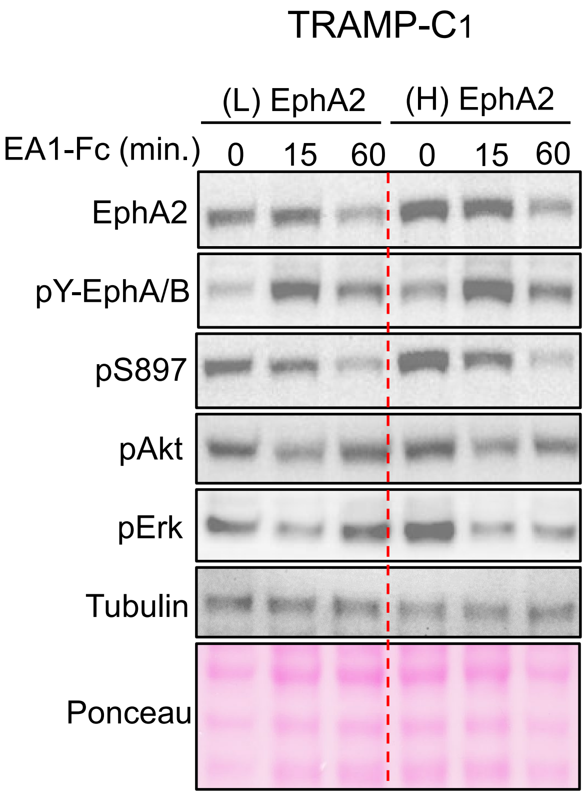

TRAMP-C1
(L) EphA2
(H) EphA2
EA1-Fc (min.)
0
15
60
0
15
60
EphA2
pY-EphA/B
pS897
pAkt
pErk
Tubulin
Ponceau


B

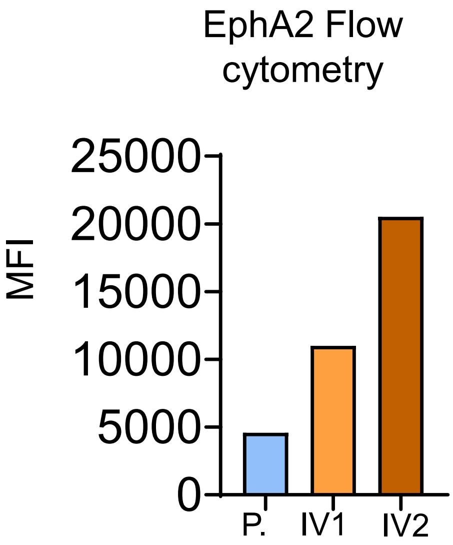

EphA2 Flow cytometry
MFI
25000
20000
15000
10000
5000
0
P.
IV1
IV2

**Fig. S2. TRAMP-C1 cells expressing higher levels of EphA2 exhibited elevated activation of Akt and ERK. A)** TRAMPC1 cells were sorted for high (H) EphA2 expression and low (L) EphA2 expression by staining with EphA2-PE and using FACS to collect the brightest 25% of cells and the lowest 25% of cells. Resultant High and Low EphA2 expressing cells were stimulated *in vitro* with dimeric EA1-Fc ligand. **B)** MPC3 cells were passaged serially in the subcutaneous space of C57Bl/6 mice as described (see methods). Relative EphA2 cell surface expression was determined by measuring mean fluorescence intensity (MFI) via flow cytometry.

Fig. S3

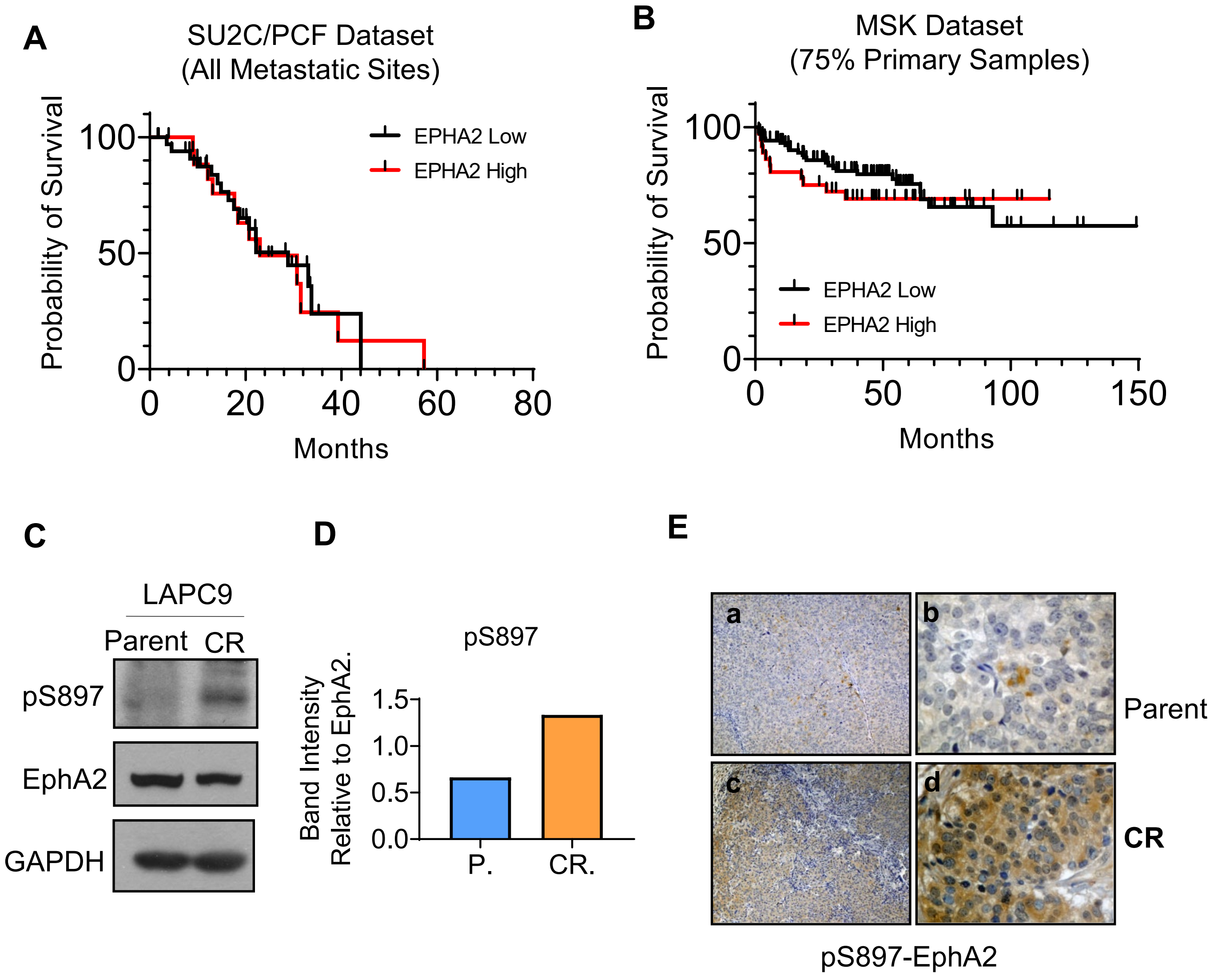
A
SU2C/PCF Dataset
(All Metastatic Sites)
Probability of Survival
100
50
0
EPHA2 Low
EPHA2 High
0
20
40
60
80
Months
B
MSK Dataset
(75% Primary Samples)
Probability of Survival
100
50
0
EPHA2 Low
EPHA2 High
0
50
100
150
Months
C
LAPC9
Parent
CR
pS897
EphA2
GAPDH
D
pS897
Band Intensity
Relative to EphA2.
1.5
1.0
0.5
0.0
P.
CR.
E
a
b
Parent
c
d
CR
pS897-EphA2

**Fig. S3. A)** Kaplan-Meier curve of patients from the SU2C/PCF dataset based on EphA2 mRNA expression considering all metastatic sites together. **B)** Survival data from the MSK dataset according to EphA2 expression. This dataset differs from the SU2C/PCF dataset in that it consist mostly (75.2%) of primary prostate tumor samples. **C)** Lysates from LAPC9 xenografts that are castration resistant (CR) androgen dependent (AD) were probed for EphA2 and pS897-EphA2. **D)** Immunoblot band quantification of pS897 signal relative to EphA2. **E)** FFPE sections from AD and CR LAPC9 xenografts were subject to IHC staining for pS897-EphA2.

### Fig. S4.

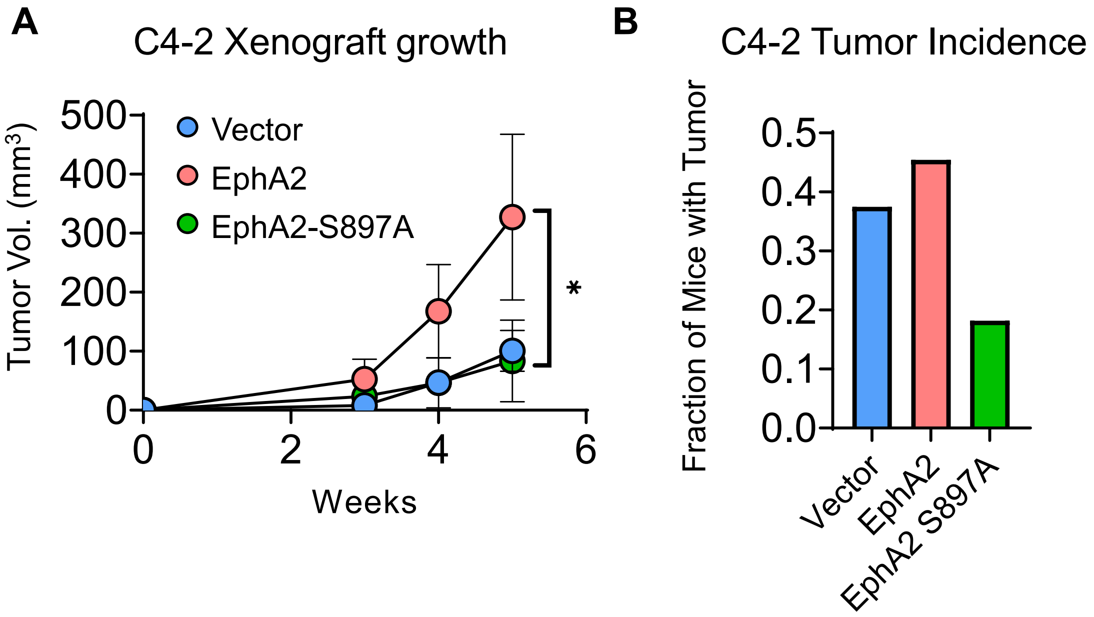


**Fig. S4. S897A mutation suppressed C4-2 xenograft tumor development. A)** C4-2 PCa cells expressing a vector control, WT-EphA2, or the S897A-EphA2 mutant were implanted subcutaneously into nude mice. **B)** C4-2 cells exhibited high latency to tumor formation with some injection sites not forming measurable tumors. This allowed for tumor incidence to be reported as the number of tumors formed relative to the total number of sites injected. Vector n = 22 sites, WT-EphA2 n = 12 sites, EphA2-S897A n = 12 sites. Unpaired t-test used for comparison of vector and WT-EphA2 at 6 weeks ($p = 0.0419$).

### Fig. S5

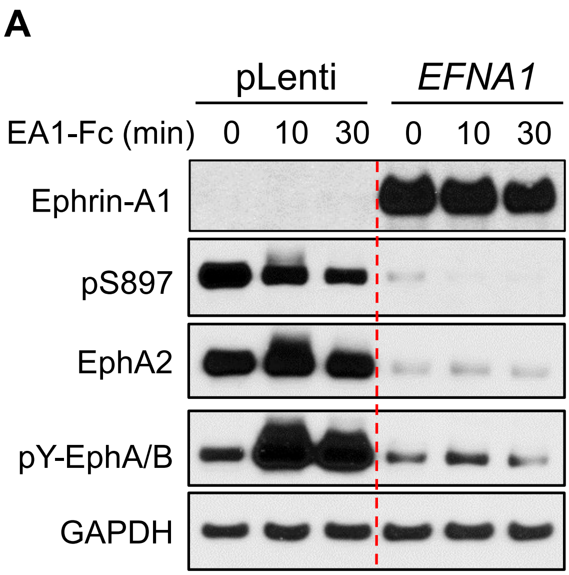


**Fig. S5. Ligand Stimulation of PC3 cells with Restored Expression of EFNA1 show Loss of EphA2-S897 Phosphorylation A)** Stimulation of Vec. and EA1 expressing PC3 cells with EA1-Fc ligand *in vitro* for 0, 10, and 30 minutes results in reduced EphA2 expression, and loss of EphA2-S897 phosphorylation signal.
